# When Does Reaction Norm GWAS Discover Plasticity? The Residual Channel in Environmental Index-Based Genetic Dissection

**DOI:** 10.64898/2026.07.27.741005

**Authors:** Shawn D. Jenkins, George L. Graef

## Abstract

CERIS-JGRA replaces the environmental mean in Finlay-Wilkinson regression with a climate-derived index, so that reaction norm association mapping can dissect the genetics of phenotypic plasticity. Its defining design choice forecloses that objective. An index correlated at *ρ* with the environmental means decomposes exactly into *ρ* times the mean plus a residual, and its loading on informative variation orthogonal to the mean, *τ*, cannot exceed √(1 − *ρ*²). Sensitivity independent of mean performance is orthogonal to the mean by definition, reaching the slope only through *τ*, and the non-centrality of a plasticity-specific test goes as *τ*². The algorithm maximizes *ρ*, minimizing by construction the channel carrying the signal it is used to detect. Simulation confirms this. A parameter-free expression reproduces observed power across 189 conditions to a root mean square error of 0.030. Power to detect plasticity-specific loci falls from 0.64 at *ρ* = 0 to 0.009 at *ρ* = 0.99 and 0.003 at *ρ* = 0.996, invariant to architecture, coupling and environment count. Holding *ρ* at 0.5 while reducing *τ* from 0.87 to zero drops power from 0.576 to 0.000: *ρ* governs discovery as a proxy, not a cause. Panel size compensates, scaling as 1 / (1 − *ρ*²): 125-fold at *ρ* = 0.996. Reimplemented on published sorghum data, the search returns the published index, photothermal time 18 to 43 days after planting, first of 27,144 candidates, at *τ* = 0.083. We recommend reporting *τ* beside *ρ*, and reading slope loci found below *τ* ≈ 0.15 as mean-performance loci.

**Article summary:** Crop geneticists commonly replace a field trial’s average yield with an index built from weather data, then map the genes controlling how a plant responds to its environment. We show by algebra that this cannot work as intended. The index is chosen to track the trial average as closely as possible, and the closer it tracks, the less room remains for the independent signal that plasticity genes need to be detected. Simulation and a reanalysis of published sorghum data agree. The search that makes an index good for predicting performance is the same search, run backwards, that makes it useless for finding genes.

## INTRODUCTION

Phenotypic plasticity, the capacity of a genotype to produce different phenotypes across environments, is a fundamental evolutionary property (Bradshaw, 1965; Des Marais et al., 2013; Schneider, 2022) and, expressed as genotype-by-environment interaction (G×E), a practical obstacle to selection (Cooper & DeLacy, 1994; Van Eeuwijk et al., 2016). Whether it is governed by dedicated regulatory loci or emerges as a by-product of the same genes controlling mean performance is among the oldest questions in quantitative genetics (Falconer, 1952; Via, 1993), and the answer sets breeding strategy: loci distinct from mean performance can be targeted independently; loci shared with it cannot. The standard framework regresses each genotype’s performance on the environmental mean (Finlay & Wilkinson, 1963). The construction is circular, because every genotype contributes to the mean against which it is regressed, and it cannot reach environments not yet sampled (Crossa, 1990). CERIS-JGRA replaces that mean with an index derived from climate data (X. Li et al., 2018, 2021): an exhaustive search over all combinations of environmental parameter and growing-period window returns the candidate whose absolute Pearson correlation with the vector of environmental means, hereafter *ρ*, is strongest. Slope and intercept estimated against that index become GWAS phenotypes, and integration with genomic information (JGRA) predicts performance in untested environments (Guo et al., 2020; He et al., 2025).

The framework has since been applied in sorghum (X. Li et al., 2018; Mu et al., 2022; Wei et al., 2025), rice (Guo et al., 2020), maize (He et al., 2025; Kusmec et al., 2017; X. Li et al., 2021; Ozair et al., 2025; Tibbs-Cortes et al., 2024), wheat (Han et al., 2025; B. Li et al., 2025), tomato (Diouf et al., 2020), soybean (Happ et al., 2021; Xavier et al., 2018), barley (Lacaze et al., 2009), rapeseed (Zeng et al., 2025), sunflower (Mangin et al., 2017), and cotton (Souaibou et al., 2025). One question remains unresolved: when *ρ* is very high, does slope GWAS identify loci controlling plasticity independently of mean performance, or does it rediscover the loci intercept GWAS already finds?

### The residual channel

High *ρ* is the explicit objective of that search, not an incidental property of the index it returns, and the consequences for genetic discovery follow from the structure of the index alone, independent of any assumption about genetic architecture. Let *y* denote the standardized vector of environmental means across *m* environments and *x* the standardized index. For any *ρ* = cor(*x*, *y*), the index decomposes exactly as *x* = *ρy* + *τz* + *νw*, where *z* is a unit-variance environmental axis orthogonal to *y*, *w* is orthogonal to both, and *τ*² + *ν*² = 1 − *ρ*². This is the ordinary regression identity and holds without approximation for any index and any *ρ*. The quantity *τ* is the loading of the index on environmental variation that is real, informative and independent of the environmental mean, and *ν* its loading on variation carrying none; every index therefore satisfies *τ* ≤ √(1 − *ρ*²), with equality only when the entire residual is informative.

For a genotype *i* with sensitivity *b_i_* to the mean axis and *s_i_* to the orthogonal axis, the Finlay-Wilkinson slope estimated against *x* is *β_i_* = *ρ*(*σ*_E_ + *b_i_*) + *τs_i_* + *ε_i_*, an exhaustive partition in which environmental sensitivity independent of mean performance, orthogonal to *y* by definition, reaches *β* only through the second term, with loading *τ*. The bound is definitional rather than empirical: whatever portion of a genotype’s environmental response is independent of mean performance can enter an index-based reaction norm slope only through that residual channel, and *ρ* caps its magnitude at √(1 − *ρ*²).

A marker with additive effect *α* on *s* has effect *τα* on *β*, so its association test carries non-centrality *λ* ∝ *n τ*² *α*² 2*f*(1 − *f*) / [*ρ*² *V*(*b*) + *τ*² *V*(*s*) + *σ*²*_ε_*], where *V*(*b*) and *V*(*s*) are the genetic variances of mean-axis and orthogonal-axis sensitivity and *σ*²*_ε_* is slope estimation error. Increasing *ρ* acts on this expression twice, in opposite terms and in the same direction. In the numerator *τ*² ≤ 1 − *ρ*², so at *ρ* = 0.99 a plasticity-specific locus retains at most 2% of the non-centrality it would carry against an index uncorrelated with the mean. In the denominator the *ρ*(*σ*_E_ + *b_i_*) term loads mean-axis sensitivity onto *β* with weight *ρ*², inflating the variance the test is scaled against. Loci that track mean performance become the noise floor for loci that do not.

The CERIS objective is to maximize *ρ*. A single optimization therefore closes the channel carrying plasticity-specific signal and inflates the variance that signal is tested against: the objective function is the reciprocal of the quantity genetic dissection requires. This reframes rather than contradicts the demonstrated value of the framework. Prediction exploits the *ρ* term, and the reported accuracies are not in question here; discovery requires the *τ* term, which the same maximization destroys. The two applications are not merely separable; they are opposed, and no choice of window reconciles them, because they are the same search conducted in opposite directions.

The constraint has a quantitative-genetic precedent and has been reached independently. Reaction norm shape cannot change independently of mean performance once the additive genetic correlation between character states approaches one (Via & Lande, 1985); the bound derived here is that statement made about the regressor rather than about the genotypes. The Finlay-Wilkinson slope has separately been shown to track the first factor loading of a factor analytic model when environments are ordered by mean yield (Sadras et al., 2026). The constraint is algebraic rather than phenotypic: in the Maize1000 subset of the Genomes to Fields hybrid panel (1,000 hybrids, seven environments), the correlation between slope and genotypic mean was near zero (*r* = 0.00-0.20) despite |*ρ*| = 0.953-0.989 for all four traits (He et al., 2025).

Here we derive the bound, confirm it by simulation with a parameter-free prediction of power, separate *ρ* from *τ* empirically to establish which of the two is causal, and apply a reimplementation of the search to the dataset in which the framework was introduced.

## MATERIALS AND METHODS

### Simulation model and QTL architecture

To formalize the relationship between *ρ* and architectural overlap, we developed a simulation framework that generates multi-environment phenotypic data under controlled genetic architectures and environmental index structures. The simulation models *n* genotypes (default 500) across *m* environments (default 12) with *p* biallelic markers (default 2,000), incorporating five QTL classes following the Des Marais *et al*. (2013) taxonomy:

i. 10 mean-only QTL affecting intercept but not slope;
ii. 10 differential sensitivity (DS) QTL where slope effects are a scaled function of intercept effects (*b*_slope_ = *c*_DS_ ⋅ *b*_intercept_, with coupling parameter *c*_DS_ = 0.8);
iii. 5 independent plasticity QTL affecting only the environmental sensitivity axis orthogonal to the mean;
iv. 3 conditional neutrality (CN) QTL active only when the environmental driver exceeds a threshold (ℎ > 0); and
v. 2 antagonistic pleiotropy (AP) QTL with coupling parameter *c*_AP_ = −0.6, producing sign-reversing effects across environments.

These proportions (i.e., DS 33%, mean-only 33%, independent 17%, CN 10%, AP 7%) reflect the empirical predominance of DS (Des Marais et al., 2013). The default coupling parameter *c_DS* = 0.8 represents the hypothesis that slope effects of DS QTL are a large fraction of their intercept effects; while Des Marais et al. (2013) established the prevalence of DS, the coupling magnitude is not directly estimable from existing meta-analyses. Experiment 4 varies *c_DS* from 0 to 1.0 to quantify the sensitivity of all headline results to this assumption (see Sensitivity analyses). Effect sizes for each QTL class are drawn from 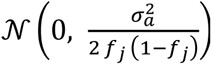, where *f_j_* is the allele frequency at marker *j*, standardizing the per-locus additive genetic variance contribution (default *σ*^2^ = 0.04 for all classes).

Genotypes are simulated under three population structure models. For the diversity panel (default), marker allele frequencies are drawn from Uniform(MAF_min_, 0.5) with MAF_min_ = 0.05, and local linkage disequilibrium is induced via AR(1) correlation blocks of 20 markers with autocorrelation *r*_AR1_ = 0.7. Diploid genotypes are constructed by summing two independently drawn haplotypes. For the biparental cross, two nearly homozygous parents are created (allele frequency 0.95 vs. 0.05) and *n* progeny are generated by simulating Poisson-distributed crossover events (rate = 0.02 per marker interval) on each gamete. For the NAM design, 10 families share a common parent crossed to 10 diverse founders (allele frequencies drawn from Uniform(MAF_min_, 0.5)), with recombination simulated independently within each family at the same rate. Only the diversity panel is reported: under a linear mixed model fitted with the genomic relationship matrix, the false positive rate for slope GWAS at *ρ* = 0.996 was 0.0008 in the diversity panel against a nominal threshold of 2.5 × 10⁻⁵, against 0.017 in the biparental population and 0.006 in the NAM population, with genomic inflation factors of 0.91, 0.55 and 1.54 respectively, so the biparental and NAM arms are not calibrated and their power estimates are not interpretable. Phenotype model. For each genotype *i* in environment *j*, the phenotypic value is generated as:

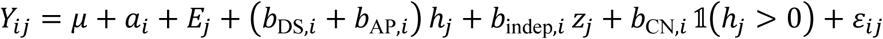

where *μ* = 50, *a_i_* is the intercept breeding value, *E_j_* = *σ_E_* ⋅ ℎ*_j_* is the environmental effect (*σ_E_* = 3.0), ℎ*_j_* are the environmental driver values drawn from N(0,1), *z_j_* is an orthogonal environmental axis constructed by residualizing an independent N(0,1) vector on ℎ and standardizing, and 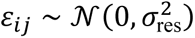 with *σ*_res_= 1.0.

The environmental index *x* is constructed to achieve a target correlation *ρ* with the vector of environmental means:

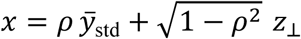

where *z*_⊥_ is orthogonal to *y̅*_std_, partitioned between the true independent environmental axis *z* (controlled by *κ*, default 1.0) and pure noise. This construction provides exact control over *ρ* while maintaining realistic distributional properties.

### Experimental design

We conducted 25 simulation experiments spanning more than 200,000 total replicates (Table 1). The primary experiment (Experiment 1) swept *ρ* from 0.00 to 0.999 over a grid of 105 values with 1,000 replicates each (105,000 replicates in total; two pairs of grid values coincide to four decimal places, leaving 103 distinct levels), establishing the core relationship between *ρ* and all overlap metrics. Experiment 2 quantified the null model (no true plasticity QTL) to establish baseline false-positive overlap. Experiments 3–13 systematically varied one factor at a time. Experiments 14–25 address additional methodological factors that could influence the generality of the findings.

**Table 1.** Summary of simulation experiments. All experiments use the default architecture (10 mean-only, 10 DS, 5 independent, 3 CN, 2 AP QTL) unless noted.

| Exp. | Name | Design | Reps/$\rho$ | Total runs |
| --- | --- | --- | --- | --- |
| 1 | $\rho$ sweep | $\rho = 0.00\text{--}0.999$ (105 levels); default architecture | 1,000 | 105,000 |
| 2 | Null model | No independent, CN, or AP QTL (DS-only) | 300 | 31,500 |
| 3 | Kappa | Index informativeness $\kappa = 0, 0.5, 1.0$ | 200 | 4,200 |
| 4 | DS coupling | Coupling c DS = 0, 0.25, 0.5, 0.75, 1.0 | 200 | 7,000 |
| 5 | Env. range | Wide vs. narrow environmental range | 200 | 2,800 |
| 6 | Env. count | n env = 3, 4, 5, 6, 8, 12, 16, 20 | 200 | 11,200 |
| 7 | Pop. structure | Diverse, biparental, NAM | 150 | 3,150 |
| 8 | Architecture | DS-only, DS+indep, DS+CN, DS+AP, full | 200 | 7,000 |
| 9 | Sample size | $n = 50, 100, 200, 500, 1000, 2000, 5000$ | 100 | 3,500 |
| 10 | Nonlinearity | Quadratic reaction norms (var quad = 0.03) | 200 | 2,800 |
| 11 | Effect size | var a indep = 0.01, 0.02, 0.04, 0.08 | 200 | 4,000 |
| 12 | Rotation test | Permutation validation (200 perms/rep) | 50 | 350 |
| 13 | Multi-index | Secondary index Z orthogonal to mean | 150 | 1,050 |
| 14 | Marker density | p = 500, 1000, 2000, 5000, 10000 | 100 | 2,500 |
| 15 | Heritability | $\sigma_{\text{res}} = 0.5\text{--}8.0$ (8 levels) | 100 | 4,000 |
| 16 | QTL count | 10–1,000 total QTL (6 configs) | 100 | 3,000 |
| 17 | Hetero. residuals | Gamma-distributed residual variance | 100 | 1,000 |
| 18 | PCA/LMM GWAS | PCA-corrected and LMM GWAS methods | 100 | 3,000 |
| 19 | Effect dist. | Equal vs. unequal effect variances | 100 | 1,000 |
| 20 | Missing data | 0%, 5%, 10%, 20% missing phenotypes | 100 | 2,000 |
| 21 | Replication | 1–4 within-environment replicates | 100 | 2,000 |
| 22 | Convergence | 10–300 replicates (diagnostic) | 300 | 900 |
| 23 | Seed sensitivity | 5 independent seed streams | 100 | 1,500 |
| 24 | LD window | Window = 0, 2, 5, 10 markers | 100 | 2,000 |
| 25 | MAF sweep | MAF min = 0.01, 0.05, 0.10, 0.20, 0.35 | 100 | 2,500 |

### Meta-analysis curation

We compiled data from 47 study-trait combinations across 27 publications spanning eleven crop species. The index-mean correlation is reported or computable for 24 of these, which are the entries listed in Table 2; the remainder report reaction norm analyses without sufficient information to recover *ρ*. Studies were included when they applied an environmental index to multi-environment trial data and performed genetic mapping for reaction norm parameters.

**Table 2.** Cross-study meta-analysis of slope-intercept architectural overlap. All 24 study-trait combinations with reported or computable *ρ*; the complete 47-entry dataset is in Supplemental Table S1. Reg. = Regime: C = Constrained (*ρ* ≥ 0.99), T = Transitional (0.85–0.99), R = Recoverable (< 0.85). N_1_ = intercept markers; N_2_ = slope markers. Overlap = QTL overlap fraction or marker Jaccard.

| Study | Species | Pop. type | N | Envs | Trait | $\rho$ | Overlap | Reg. | Tier | $\tau$ |
| --- | --- | --- | --- | --- | --- | --- | --- | --- | --- | --- |
| Kusmec 2017 | Maize | NAM | 4928 | 23 | 23 traits | 1.000 | 0.004 | C | 1 | 0.000 |
| Tibbs-Cortes 2024 | Maize | NAM | 4928 | 11 | DTA | 0.999 | 0.004 | C | 1 | 0.045 |
| Tibbs-Cortes 2024 | Maize | NAM | 4928 | 11 | DTS | 0.998 | 0.066 | C | 1 | 0.063 |
| Li 2018 | Sorghum | Biparental | 237 | 7 | FT | 0.996 | 1.000 | C | 1 | 0.089 |
| Li 2018 | Sorghum | Biparental | 237 | 7 | PH | 0.995 | 1.000 | C | 1 | 0.100 |
| Mu 2022 | Sorghum | NAM | 771 | 6 | PH | 0.995 | 1.000 | C | 1 | 0.100 |
| Guo 2020 | Rice | Biparental | 174 | 9 | FT | 0.990 | 0.800 | C | 1 | 0.141 |
| Li 2021 | Maize | Diverse | 282 | 13 | FT | 0.970 | 0.111 | T | 1 | 0.243 |
| Ozair 2025 | Maize | Hybrid | 284 | 11 | GY (PHK76) | 0.958 | 0.500 | T | 1 | 0.287 |
| Han 2025 | Wheat | Diverse | 406 | 10 | PH | 0.940 | 0.400 | T | 1 | 0.341 |
| Han 2025 | Wheat | Diverse | 406 | 10 | FT | 0.984 |  | T | 1 | 0.178 |
| Li 2021 | Wheat | Breeding | 139 | 10 | FT | 0.920 |  | T | 1 | 0.392 |
| He 2025 | Maize | Hybrid | 1000 | 7 | 4 traits | 0.953–0.989 | 0.28–0.39 | T | 1 | 0.148–0.303 |
| Zeng 2025 | Rapeseed | Diverse | 505 | 8 | Oil content | 0.84–0.98 | 0.000 | T | 1 | 0.199–0.543 |
| Wei 2025 | Sorghum | Diverse | 306 | 14 | FT | 0.850 | 0.000 | T | 1 | 0.527 |
| Wei 2025 | Sorghum | Diverse | 306 | 14 | PH | 0.850 |  | T | 2 | 0.527 |
| Diouf 2020 | Tomato | Diverse | 156 | 4 | 10 traits | 1.000 | 0.350 | C | 1 | 0.000 |
| Lacaze 2009 | Barley | Diverse | 224 | 8 | Multiple | 1.000 | 1.000 | C | 1 | 0.000 |
| Jin 2023 | Maize | Diverse | 1404 | 5 | 23 traits | 1.000 | 0.128 | C | 1 | 0.000 |
| Happ 2021 | Soybean | Diverse | 1516 | 4 | GY | 1.000 | 0.000 | C | 1 | 0.000 |
| Ozair 2025 | Maize | Hybrid | 326 | 14 | GY (PHP02) | 0.693 |  | R | 1 | 0.721 |
| Ozair 2025 | Maize | Hybrid | 285 | 9 | GY (PHZ51) | 0.733 |  | R | 1 | 0.680 |
| Li 2021 | Wheat | Breeding | 139 | 10 | GY | 0.540 |  | R | 2 | 0.842 |
| Mangin 2017 | Sunflower | Hybrid | 317 | 14 | Oil yield | 1.000 |  | C | 2 | 0.000 |

### Empirical validation

Simulation predictions were validated against the 19-trait analysis of the maize NAM population by Tibbs-Cortes et al. (2024). For each trait, we extracted *ρ*, marker counts, gene counts, and genome-wide −log₁₀(*P*) correlations, then compared each metric to the simulation prediction at the matched *ρ* value. Cross-study validation compared simulation-predicted overlap to empirical overlap fractions across all Tier 1 entries with quantitative data. Bootstrap confidence intervals (2,000 iterations) were computed within each *ρ* level for all key metrics (Figure S1).

### GWAS model and significance thresholds

For each simulated replicate, we performed single-marker GWAS by regressing intercept and slope breeding values on centered marker genotypes under a simple linear model (*y* ∼ *X_j_* for each marker *j*). Test statistics follow a *t*-distribution with *n* − 2 degrees of freedom. Significance was determined using a Bonferroni-corrected threshold (*α* = 0.05/*p*, where *p* is the number of markers). FDR correction (Benjamini–Hochberg at *q* = 0.05) was computed in parallel to verify that findings were not artifacts of the multiple-testing procedure (Figure S3). For population-structure-aware analyses (Experiment 18), we implemented PCA-corrected GWAS (residualizing phenotypes and genotypes on the first 3 principal components of the marker matrix) and linear mixed model GWAS (using Cholesky rotation of the genomic relationship matrix K = XX′/*p* with regularization *λ* = *n*/*p*) (Figure S8). Genomic inflation factors (*λ*_GC_) were computed from the median *t*-statistic at null markers to verify calibration (Figure S4). Genomic prediction accuracy was estimated as the Pearson correlation between predicted and true breeding values under five-fold cross-validation using ridge regression BLUP (K-BLUP with *λ* = *n*/*p*), averaged across folds.

### Overlap and redundancy metrics

Architectural overlap was quantified as the proportion of Bonferroni-significant slope markers that were also Bonferroni-significant for intercept. The metric is directional by design: the question is whether slope GWAS discovers loci beyond those intercept GWAS identifies, so we ask what fraction of slope hits are redundant with intercept hits. All figures and headline values report the exact marker-level intersection without windowing; at *ρ* = 0.996 it averaged 0.61 [95% CI 0.59–0.62], against 0.46 [0.13–0.78] for the reverse direction, whose wider interval reflects replicate-level variation in the number of intercept hits. A windowed variant, genome-wide signal similarity, and detection power by QTL class are defined in Methods S1.

The standardized expected selection response for slope (*R_b_*) was computed as:

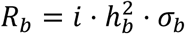

where *i* = 1.755 (selection intensity for the top 10%), 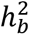 is the narrow-sense heritability of slope, and *σ_b_* is the additive genetic standard deviation of slope. *R_b_* was decomposed into components attributable to GWAS-detected markers (*R_b_*_|gwas_) and indirect selection on intercept (*R_b_*_|ind_) to quantify the practical breeding value of slope GWAS hits. The Via–Lande genetic correlation (*r_G_*) between character states was estimated by projecting the genotype-by-environment covariance matrix onto the reaction norm basis (intercept and slope of x), following the matrix formulation of Lande (Lande, 1979) (Figure S2). AMMI-1 decomposition was computed for each replicate to estimate the correlation between FW slope and the first interaction principal component (IPCA1), providing a model-free measure of the dimensionality of G×E (Figure S7).

## RESULTS

### The residual channel and the loss of plasticity-specific power

Power to detect plasticity-specific QTL declined monotonically with *ρ*, from 0.641 against an index uncorrelated with the environmental mean to 0.003 at *ρ* = 0.996 (Figure 1). The decline tracked *τ* = √(1 − *ρ*²), which falls from 1.000 to 0.089 across the same range. At *ρ* = 0.95, where *τ* = 0.312, power was 0.110; at *ρ* = 0.99, where *τ* = 0.141, power was 0.009. Power to detect differentially sensitive QTL moved in the opposite direction over the same interval, rising from 0.002 to 0.463. This is the denominator term of the non-centrality expression made visible: mean-axis sensitivity loads onto the slope as *ρ* increases, and the loci that track mean performance become the variance against which plasticity-specific loci are tested.

**Figure 1.**
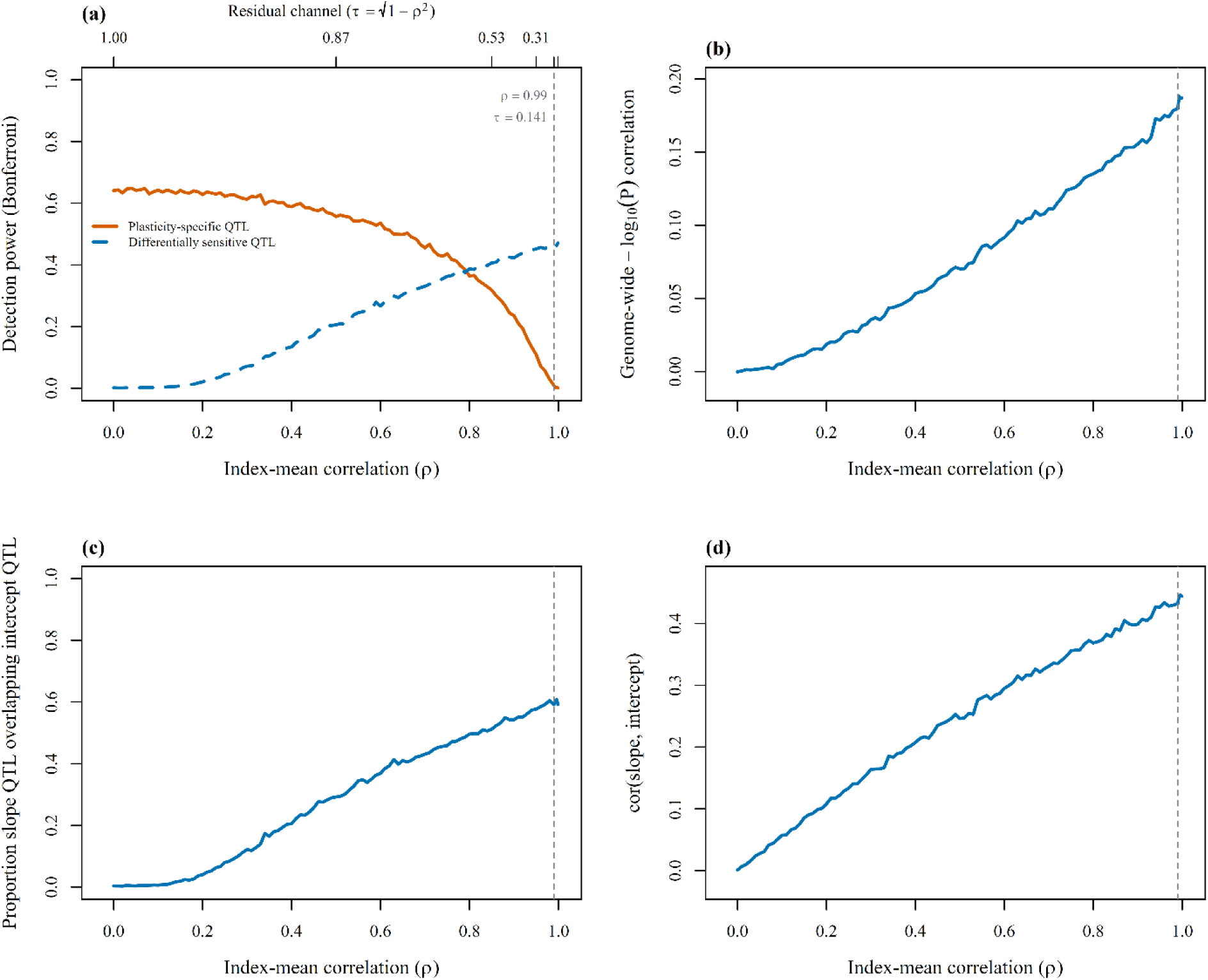
The residual channel across the *ρ* gradient (Experiment 1; 103 levels, 105,000 replicates). (a) Power to detect plasticity-specific QTL (orange, solid) and differentially sensitive QTL (blue, dashed) against *ρ*, with *τ* = √(1 − *ρ*²) on the upper axis; vertical line at *ρ* = 0.99, where *τ* = 0.141. (b) Genome-wide −log₁₀(*P*) correlation between slope and intercept GWAS. (c) QTL overlap proportion between slope and intercept GWAS. (d) Slope-intercept effect size correlation.

### ρ governs discovery as a proxy; τ is the mechanism

Under the operating assumption that the index residual is fully informative (*κ* = 1), *τ* is determined by *ρ* and the two cannot be distinguished. Varying *κ* separates them. Holding *ρ* = 0.5 and reducing *τ* from 0.866 to zero reduced power from 0.576 to 0.000. At *ρ* = 0, power was 0.656 when *τ* = 1 and 0.000 when *τ* = 0. Power at *ρ* = 0 is therefore not a property of *ρ*; it is a property of *κ*.

The denominator term separates in the same design. Holding *τ* at approximately 0.25 and increasing *ρ* from 0 to 0.95 reduced power from 0.273 to 0.043, a sixfold decline at constant signal loading. Across the full 35-cell grid, power correlated with *τ* at *r* = 0.958 and with *ρ* at *r* = −0.672; restricting to the 28 cells in which the index residual carries information (*κ* > 0) gives 0.957 and −0.810 (Figure 2).

**Figure 2.**
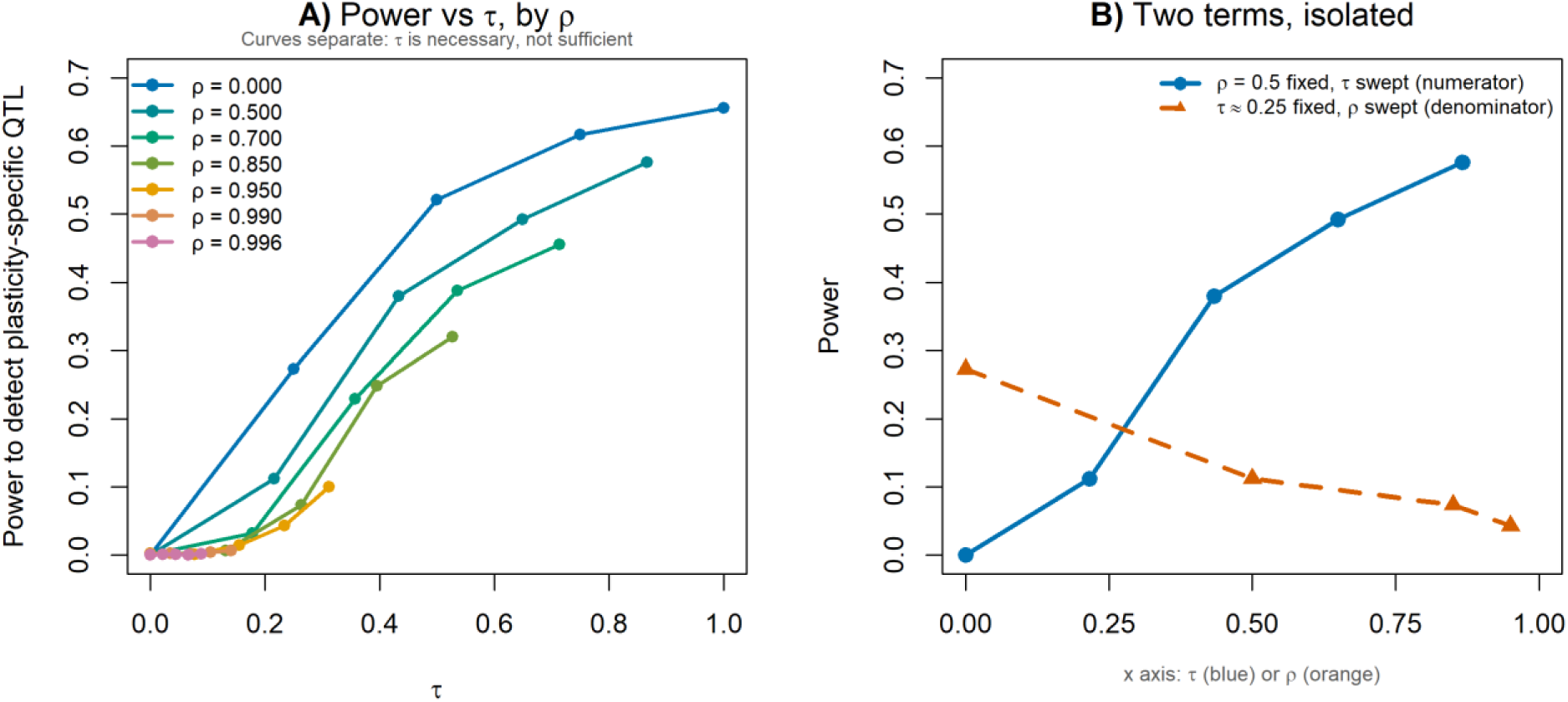
*ρ* is a proxy; *τ* is the mechanism (Experiment 3; 7 *ρ* × 5 *κ* levels, 200 replicates per cell). (a) Power to detect plasticity-specific QTL against *τ*, one line per *ρ* level; the curves separate, so *τ* is necessary but not sufficient. (b) The two terms of the non-centrality isolated. Blue: *ρ* held at 0.5 while *τ* is swept, varying the numerator alone, and power moves from 0.576 to 0.000. Orange: *τ* held at approximately 0.25 while *ρ* is swept, varying the denominator alone, and power moves from 0.273 to 0.043.

### A parameter-free prediction of power

Substituting the measured Var(*β*^) for its modeled counterpart yields a prediction of power with no free parameters: *λ* = *n τ*² *σ*²_a_ / Var(*β*^), in which *n*, *τ* and *σ*²_a_ are fixed by the design and Var(*β*^) is observed. Because per-locus effects are drawn from a normal distribution, the quantity *α*² 2*f*(1 − *f*) is distributed as *σ*²_a_ *χ*²₁, and power is the expectation of the non-central chi-square tail probability over that distribution rather than its value at the mean.

Across 189 conditions in which *ρ*, *κ*, panel size and environment count were varied independently, and spanning *λ* from 0.01 to 376, predicted and observed power agreed to a root mean square error of 0.030 (*r* = 0.995; Figure 3). The mean residual was below 0.006 at every *ρ* ≥ 0.85, with a maximum absolute error of 0.034 across those cells. The prediction was conservative at low *ρ*, understating power by up to 0.13 at *ρ* = 0, where the variance approximation is weakest. No parameter was estimated from the outcome.

**Figure 3.**
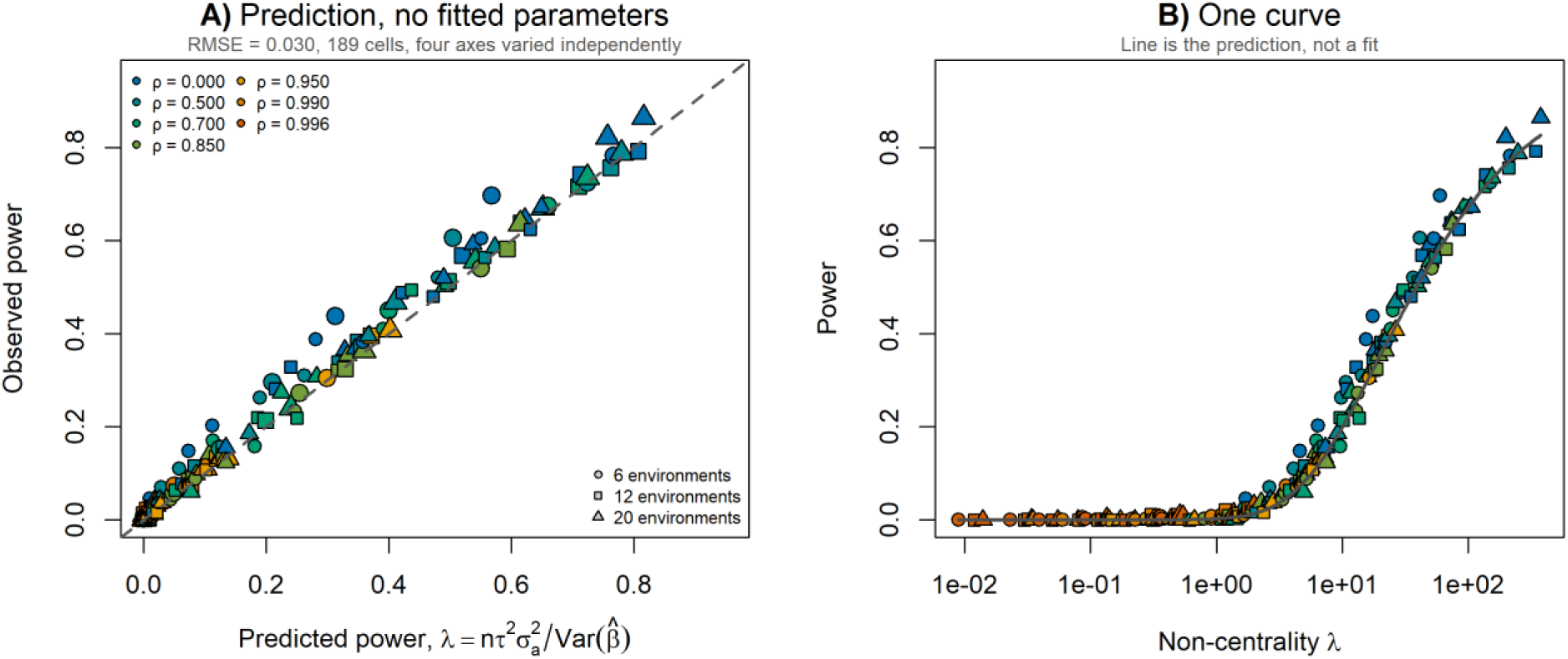
A parameter-free prediction of power (189 conditions; *ρ*, *κ*, panel size and environment count varied independently; 100 replicates per cell). (a) Observed against predicted power, where *λ* = n *τ*² *σ*²_a_ / Var(*β*^) and Var(*β*^) is measured rather than modeled; the dashed line is the identity, not a fit. Fill indicates *ρ*, shape indicates environment count, point size indicates panel size. RMSE = 0.030. (b) Panel size required to recover a plasticity-specific locus, relative to an index uncorrelated with the environmental mean, scaling as 1 / (1 − *ρ*²).

### Invariance to architecture, coupling, environments and panel size

At *ρ* = 0.996, power to detect plasticity-specific QTL ranged from 0.001 to 0.005 across five architectures spanning a fourfold range in phenotypic slope-mean correlation (0.113 to 0.454) and a twofold range in QTL overlap (0.293 to 0.594). At *ρ* = 0.95 the same five architectures gave 0.071 to 0.247. The constraint is architecture-invariant at *ρ* ≥ 0.99 and architecture-dependent at *ρ* = 0.95, which places the operational boundary at 0.99 rather than at 0.95.

At *ρ* = 0.996, power ranged from 0.001 to 0.005 across the full range of differential-sensitivity coupling, including c_DS_ = 0, where no differentially sensitive QTL exist and power to detect them is exactly zero. The default coupling of 0.8 does not generate the result, and no assumption about the prevalence or strength of allelic sensitivity is required to obtain it.

Panel size compensates, because *λ* is proportional to *nτ*². At *ρ* = 0.996, power rose from 0.000 at *n* = 500 to 0.106 at *n* = 5,000; at *ρ* = 0.95 it reached 0.590 at *n* = 5,000 (Figure S9). The panel required to recover a plasticity-specific locus therefore scales as 1 / (1 − *ρ*²), which is tenfold at *ρ* = 0.95, 50-fold at *ρ* = 0.99 and 125-fold at *ρ* = 0.996 relative to an index uncorrelated with the mean. Environment count does not compensate. Power at *ρ* = 0.996 remained between 0.000 and 0.004 from three to twenty environments, because the number of environments enters *λ* only through the estimation error term (Figure S10).

### Prediction and discovery diverge

Prediction and discovery draw on different terms of the same decomposition, and the simulation separates them. Genomic prediction accuracy for the intercept was stable across the *ρ* gradient, while accuracy for the slope rose with *ρ*: the index becomes a better predictor of performance precisely as it becomes a worse instrument for dissection. This is the *ρ* term doing the work that the *τ* term cannot. It is also why the framework’s reported prediction accuracies are not in question here and are not evidence that slope GWAS is recovering plasticity (Figure 4).

**Figure 4.**
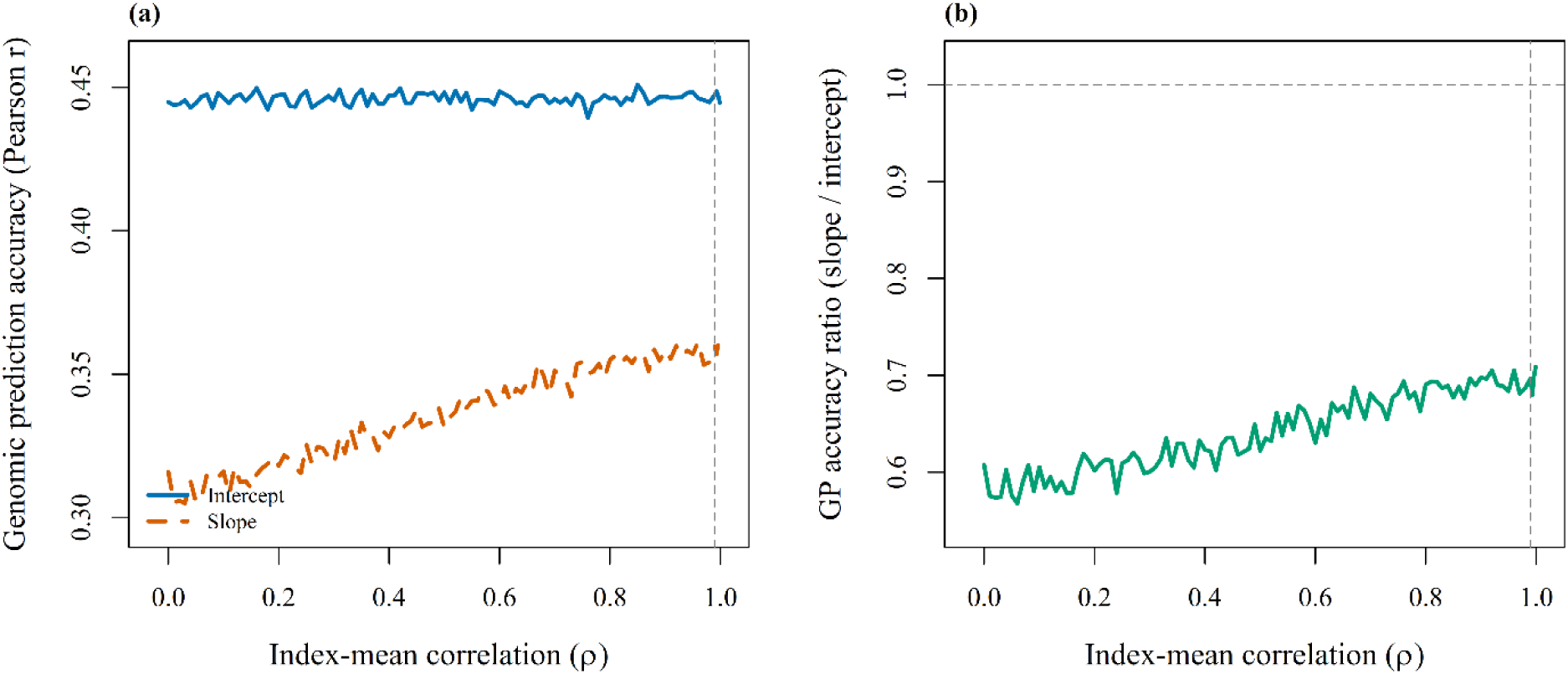
Genomic prediction accuracy across the *ρ* gradient (Experiment 1). (a) Prediction accuracy (Pearson *r*, five-fold cross-validation) for intercept (blue, solid) and slope (orange, dashed). (b) GP accuracy ratio (slope/intercept); dashed line at 1.0 for reference.

### Application to the sorghum data of Li et al. (2018)

We reimplemented the CERIS search against the data distributed by the authors: 237 recombinant inbred lines evaluated in seven environments, with daily values for seven environmental parameters over 122 days after planting. Enumerating every parameter-window combination with *j* > *i* + 5 across the four parameters named in the published description yielded 27,144 candidates. The search recovered photothermal time from 18 to 43 days after planting as the highest-ranked candidate at *ρ* = 0.9965, which is the index reported by Li et al. (2018).

That index carries *τ* = 0.083. Under the bound derived above, a plasticity-specific locus in this population retains at most 0.7% of the non-centrality it would carry against an index uncorrelated with the environmental mean, and the panel required to recover such a locus at the power available when *τ* = 1 is 145 times the 237 lines evaluated. The constraint is not an artifact of the simulation; it is a property of the index selected in the study that introduced the framework.

### Where the published literature sits

To establish where the published literature sits on the *ρ* axis, we compiled 47 study-trait combinations across 27 publications spanning eleven crop species, of which 24 report or permit calculation of *ρ* (Table 2). Twelve of those 24 lie at *ρ* ≥ 0.99, where *τ* ≤ 0.141 and no architecture we examined retained power to detect plasticity-specific loci. Flowering time, the trait most extensively analyzed with this framework, is at or above 0.99 in every entry. The constraint is not a hypothetical regime; it is where the method is most often applied.

Overlap is reported on incommensurable scales across these studies: QTL colocalization in small biparental populations, marker-level Jaccard indices in large mapping panels, and qualitative gene-list comparisons. Predicted and observed overlap therefore correlate only weakly across all entries (*r* = 0.22). Within the one class the simulation maps onto directly (QTL colocalization in small biparental studies), four of the six entries at *ρ* > 0.99 show at least 80% colocalization.

## DISCUSSION

### The prediction-discovery disconnect

Prediction accuracy and genetic discovery are separable properties of the CERIS-JGRA framework, and only one of them is in question here. The prediction applications are well supported: He et al. (2025) report accuracies of 0.619 to 0.975 across traits, and Ozair et al. (2025) found photothermal-ratio models comparable to full G×E models while running 192 times faster. Our simulation shows why. Genomic prediction accuracy for slope rises with *ρ* (GP ratio 0.61 at *ρ* = 0 to 0.70 at *ρ* = 0.996), not because plasticity loci are better captured, but because slope absorbs more heritable intercept signal through DS coupling. Prediction succeeds by exploiting shared architecture; a discovery claim requires demonstrating distinct architecture. The reported accuracies are therefore not evidence that slope GWAS recovers plasticity, and nothing here disputes them.

### Breeding consequences and the location of the boundary

The breeding consequence is direct. Marker-assisted selection on slope hits loses roughly 40% of its effectiveness between *ρ* = 0 and *ρ* = 0.996, with R_b|gwas_ declining from 1.11 to 0.65, because at high *ρ* those hits increasingly tag differentially sensitive effects that improve the intercept more than the independent component of plasticity. This is not a failure of GWAS methodology; it is a consequence of the architecture. A breeder who wants to shift environmental sensitivity specifically needs loci with independent slope effects, and at high *ρ* those loci are statistically undetectable. The boundary at *ρ* = 0.99 is a functional threshold on a smooth, monotonic curve (Figure 1a), not a natural discontinuity: it is located where the constraint stops depending on genetic architecture. At *ρ* = 0.95, power ranges from 0.071 to 0.247 across the five architectures examined, a 3.5-fold spread, so the outcome there depends on assumptions a practitioner cannot check; at *ρ* = 0.99 the same architectures give 0.005 to 0.027. Earlier drafts placed the boundary at 0.95 on the basis of the overlap metric; the power decomposition does not support that placement, and because *τ* = √(1 − *ρ*²) is smooth, no threshold is sharp.

### Diagnostic criteria and alternatives

Three criteria follow. First, report *ρ*, and report *τ* = √(1 − *ρ*²) beside it: *ρ* is computable in every study that uses an index, and *τ* follows from it by arithmetic. Below *τ* ≈ 0.15, corresponding to *ρ* 0.99, index-based slope GWAS identified no plasticity-specific loci at any architecture, coupling strength or population structure we examined, and the panel required to do so scales as 1 / (1 − *ρ*²): tenfold at *ρ* = 0.95, 50-fold at *ρ* = 0.99 and 125-fold at *ρ* = 0.996. Slope QTL recovered in that regime should be interpreted as mean-performance QTL whose environmental sensitivity is a consequence of their effect size. We offer no overlap benchmark: at fixed *ρ* = 0.996 overlap ranges from 0.29 to 0.59 across architectures, ten times the replicate confidence interval, and a practitioner does not know their architecture in advance. In its place, the rotation test offers a per-dataset check: it compares the observed count of unique slope QTL against a pure differential-sensitivity null, rejecting that null at low ρ and losing power to do so as ρ approaches one, where the rejection rate falls to the nominal 0.05 (Experiment 12; Figure S5).

Second, assess environmental range and nonlinearity. Reliable slope estimation requires at least four environments spanning 38% of the full environmental range (Guo et al., 2024), and our Experiment 6 finds the −log₁₀(*P*) correlation unstable below six.

Third, test for multi-dimensional G×E structure. Where the optimal environmental index differs across genetic backgrounds (Ozair et al., 2025), or several environmental parameters each explain unique slope genetic variance (He et al., 2025; Millet et al., 2019), a single index captures one dimension of the G×E landscape and misses the rest: fitting two orthogonal indices simultaneously resolved 1.8× more unique slope QTL than either alone at *ρ* > 0.90 (Figure S6).

Where the criteria fail, alternatives exist that do not reduce the environmental space to a single axis. The envGWAS framework (Monteverde et al., 2019) tests marker-environment associations directly rather than through a two-stage slope/intercept decomposition, side-stepping the *ρ* constraint entirely. Kernel-based approaches (Jarquín et al., 2014) model G×E through environmental covariance matrices that preserve the multi-dimensional structure of the environmental space, and remain informative because the genetic covariance matrix is not singular even at *ρ* = 0.996, where it reaches only *rG* ≈ 0.45. Multi-index reaction norm models decompose G×E into distinct biological axes, as Zeng et al. (2025) demonstrated in rapeseed, where three independent indices yielded near-zero intercept-slope correlation and identified five plasticity-specific genes. And when CERIS returns a window with *ρ* below 0.85, the residual channel reopens: slope GWAS gains real independence and should be pursued.

The CERIS-JGRA framework was designed to bring environmental specificity to reaction norm analysis, and its predictive utility has been demonstrated across species (He et al., 2025; Ozair et al., 2025). Our analysis shows that when the environmental index achieves very high correlation with the environmental mean, that specificity is algebraic rather than biological, and slope GWAS recovers what mean GWAS already reveals.

### Limitations

Our genetic model is purely additive; epistatic interactions between mean and plasticity loci could create additional independence or redundancy depending on their sign and structure, and we cannot quantify that without an explicit epistatic simulation. The default simulation (500 genotypes, 2,000 markers) is smaller than typical crop GWAS panels, though relative patterns were stable across 50 to 5,000 genotypes and 500 to 10,000 markers (Experiments 9 and 14). The model also assumes a universal environmental gradient, in which all genotypes respond to the same index and differ only in slope and intercept; Ozair et al. (2025) showed that the optimal window varies by genotype for grain yield, so our analysis is most applicable where that assumption approximately holds, as it does for photoperiod-driven flowering time and likely does not for traits shaped by genotype-specific stress thresholds.

The simulation reproduces the qualitative pattern in empirical data but not its trait-specific magnitude: against the traits of Tibbs-Cortes et al. (2024), observed genome-wide −log₁₀(*P*) correlations span 0.02 to 0.21 where the simulation predicts 0.17 to 0.19 at matched *ρ*, giving empirical-to-simulation ratios of 0.12 to 1.17 (Supplemental Table S2). Those authors report more intercept than slope markers in 12 of their 19 traits, which agrees in direction with the excess predicted here (1.26 to 1.46 across *ρ* = 0.5 to 0.996); the ten traits in Supplemental Table S2 include all seven of their slope-dominant traits, so that subset is not representative of the direction and we do not read it as evidence either way. Marker-level Jaccard is at or below 0.07 across those ten, where the simulation predicts true QTL overlap of 0.58 to 0.61, which is the same mechanism seen from the other side: differentially sensitive QTL contribute different effect sizes to slope and intercept, so different markers cross threshold for each, and discordance among top hits coexists with shared architecture. The residual quantitative gap is most plausibly scale, since roughly 4,928 lines and more than 20 million markers were analyzed there against 500 genotypes and 2,000 markers in 20-marker linkage blocks here.

Our meta-analysis has its own constraints. The regime distribution is lopsided: 12 of the 24 classifiable study-trait combinations fall in the Constrained regime (*ρ* ≥ 0.99), nine in the Transitional regime (0.85–0.99), and three in the Recoverable regime (*ρ* < 0.85). This reflects the field’s heavy use of flowering time and FW-like indices, which tend to produce high *ρ*, and coverage still skews toward maize (20 of 47 entries) despite the addition of rapeseed (Zeng et al., 2025), sunflower (Mangin et al., 2017), cotton (Souaibou et al., 2025), and a second large maize diversity panel (Jin et al., 2023). Several studies that would be most informative for testing the Recoverable regime do not report *ρ*, so the strongest validation applies to the regime where the conclusion is least controversial. Standardized reporting of both QTL-level colocalization and genome-wide signal correlation would enable more rigorous cross-study comparison as the meta-analytic base grows.

## DATA AVAILABILITY

Simulation code (v9, 25 experiments) and all summary data files are available at https://github.com/jenkinsshawn/ReactionNormConstraints. No new empirical data were generated; all empirical data referenced are from the original publications cited herein.

## ACKNOWLEDGMENTS

We thank the authors of the 27 studies included in the meta-analysis for making their results publicly available.

## AUTHOR CONTRIBUTIONS

S.D.J. conceived the study, developed the theory and simulations, performed the analyses, and wrote the manuscript. G.L.G. acquired funding and reviewed the manuscript.

## SUPPLEMENTAL MATERIAL

**Methods S1.** Overlap and redundancy metrics. Full definitions of the metrics summarized in Materials and Methods. Available as Methods_S1.docx in the online supplement available at https://github.com/jenkinsshawn/ReactionNormConstraints.

---

**Figure S1.**
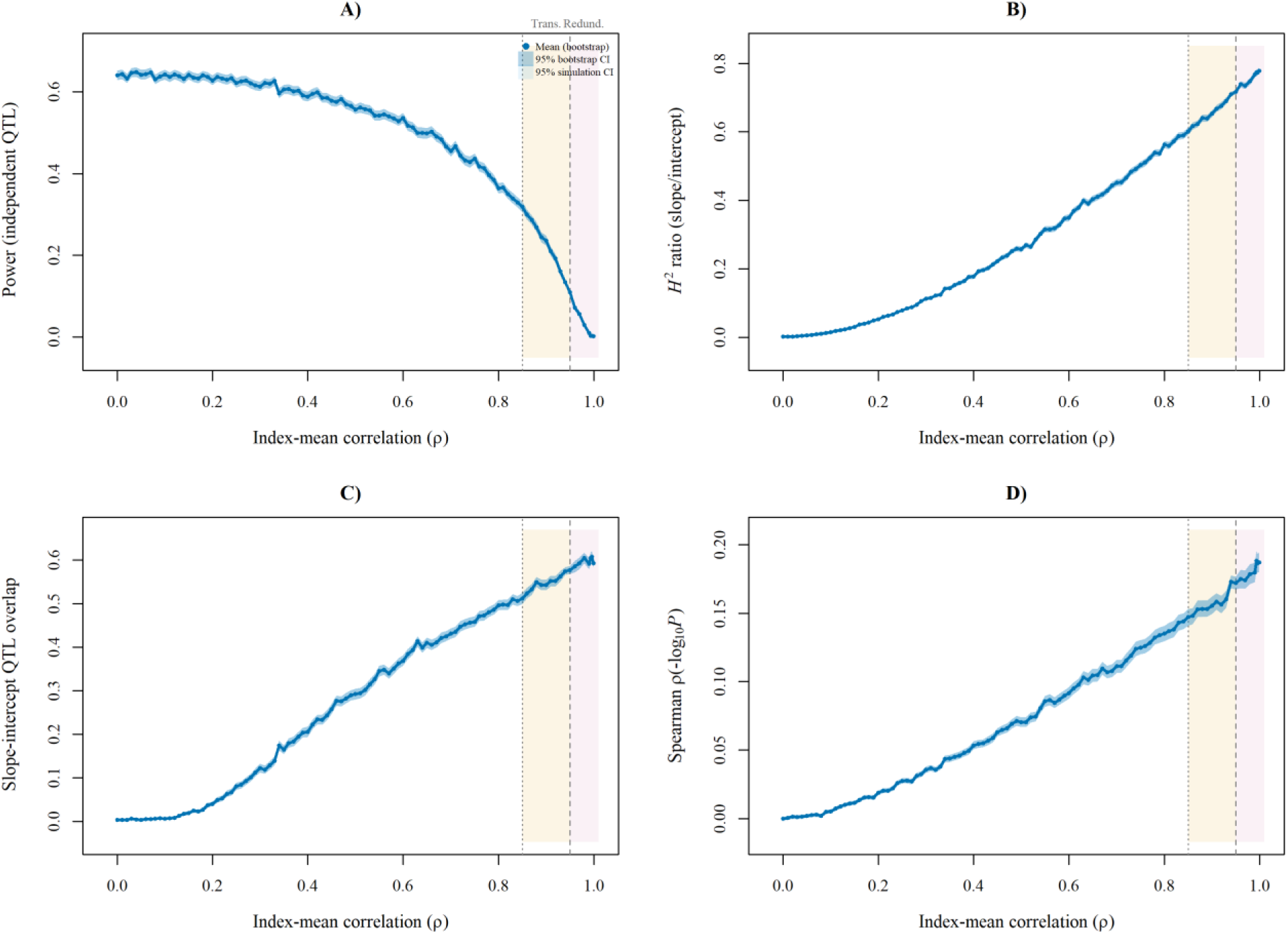
Bootstrap confidence intervals for key simulation metrics (Experiment 1; 2,000 iterations per *ρ* level). (a) Power to detect independent QTL. (b) Heritability ratio (H²slope / H²intercept). (c) Architectural overlap proportion. (d) Genome-wide P-value correlation.

**Figure S2.**
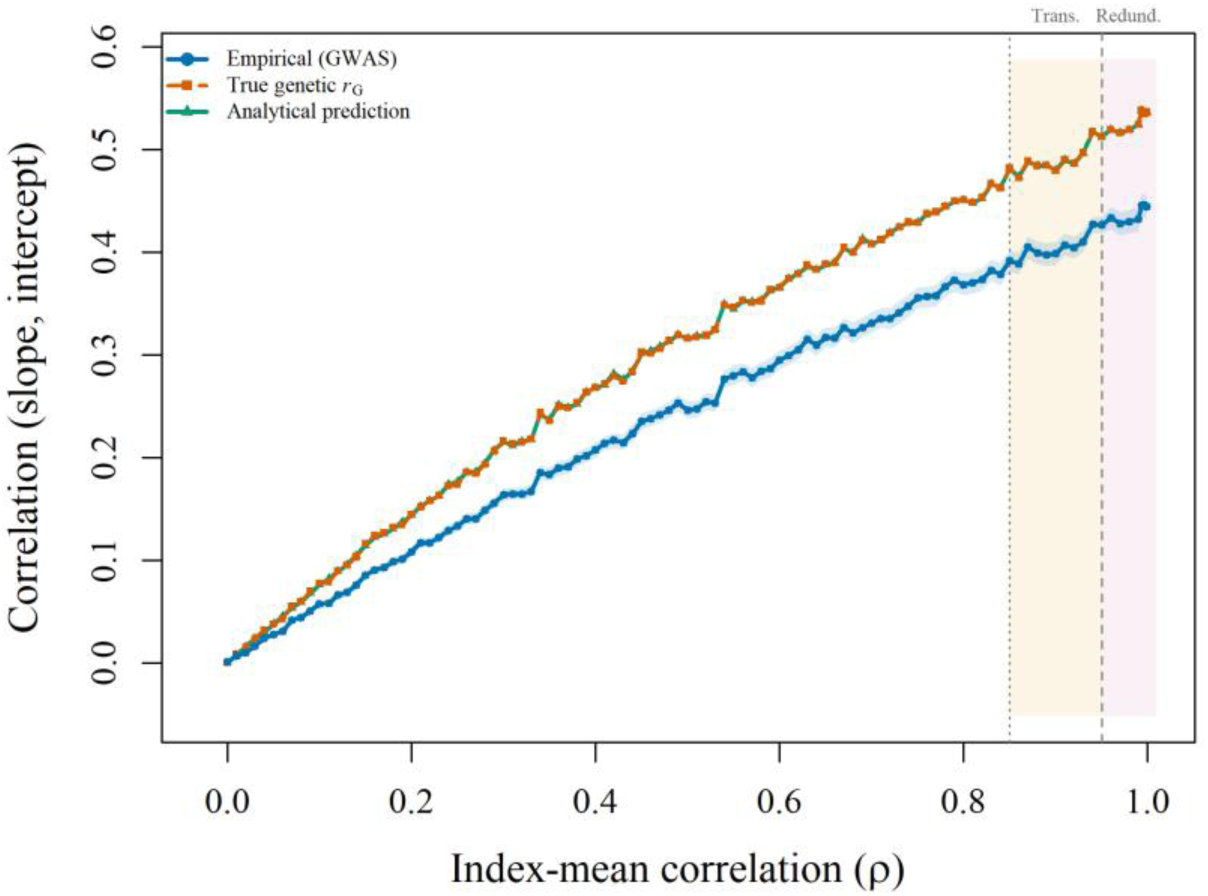
Quantitative genetics theory overlay. Correlation between slope and intercept as a function of *ρ*: empirical simulation estimate (blue, with 95% CI), true genetic correlation via the Lande (1979) estimator (orange, dashed), and Via-Lande analytical prediction (green).

**Figure S3.**
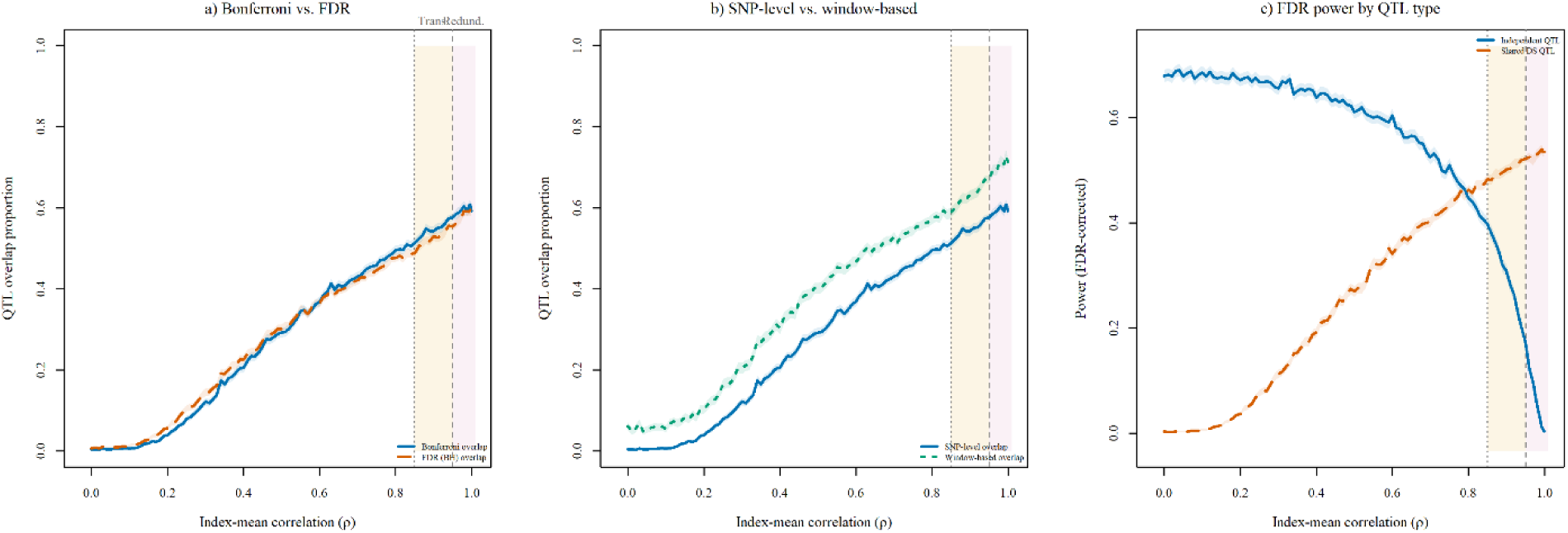
FDR robustness analysis (Experiment 1). (a) Overlap proportion under Bonferroni (blue) and FDR 5% (orange, dashed) correction. (b) SNP-level (blue) versus ±5-marker window (teal) overlap. (c) FDR-corrected power by QTL type: independent (blue) and shared DS (orange).

**Figure S4.**
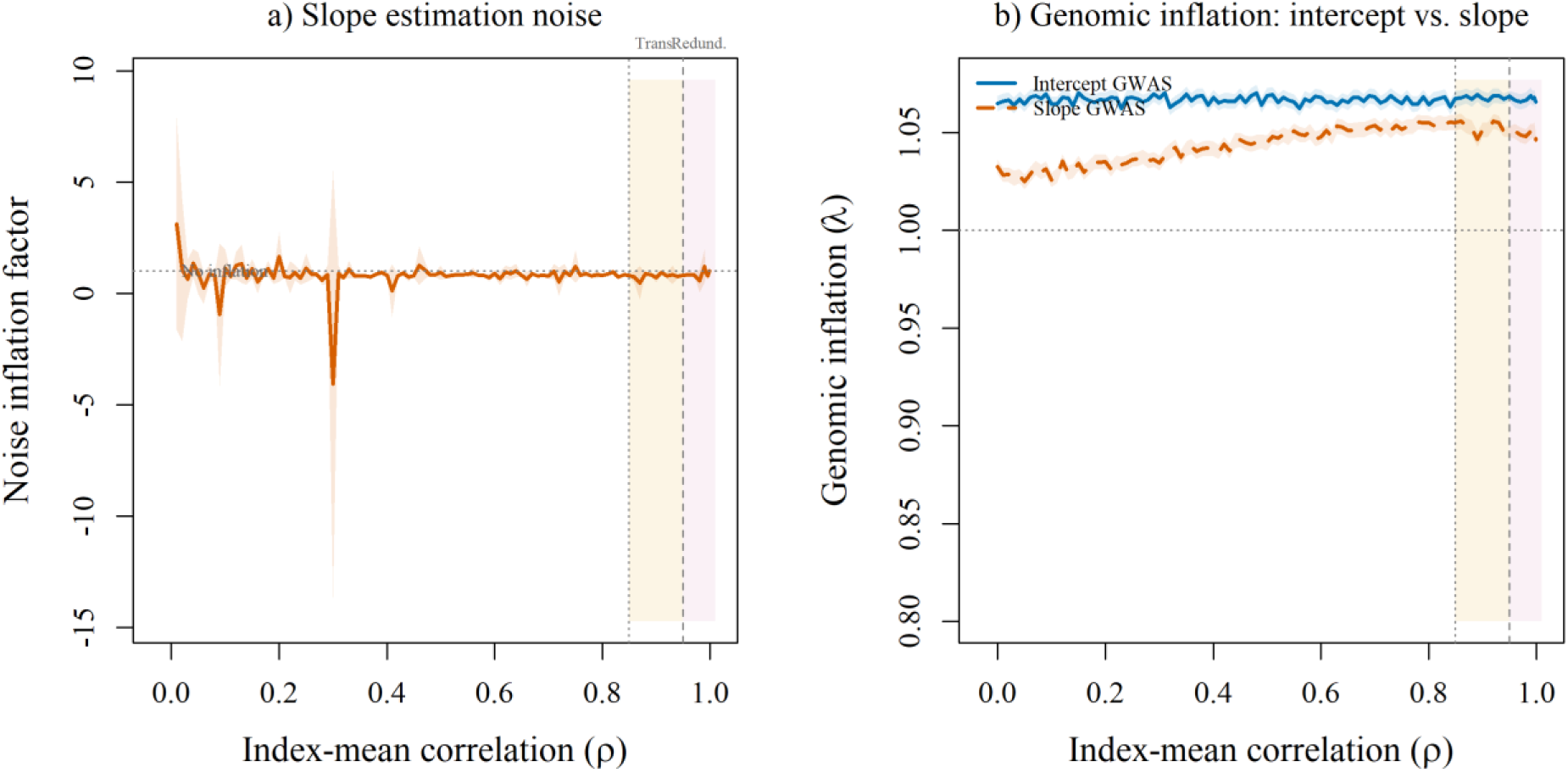
Slope estimation noise and genomic inflation (Experiment 1). (a) Noise inflation factor for slope estimation as a function of *ρ*. (b) Genomic inflation factor (*λ*_GC) for intercept (blue) and slope (orange) GWAS.

**Figure S5.**
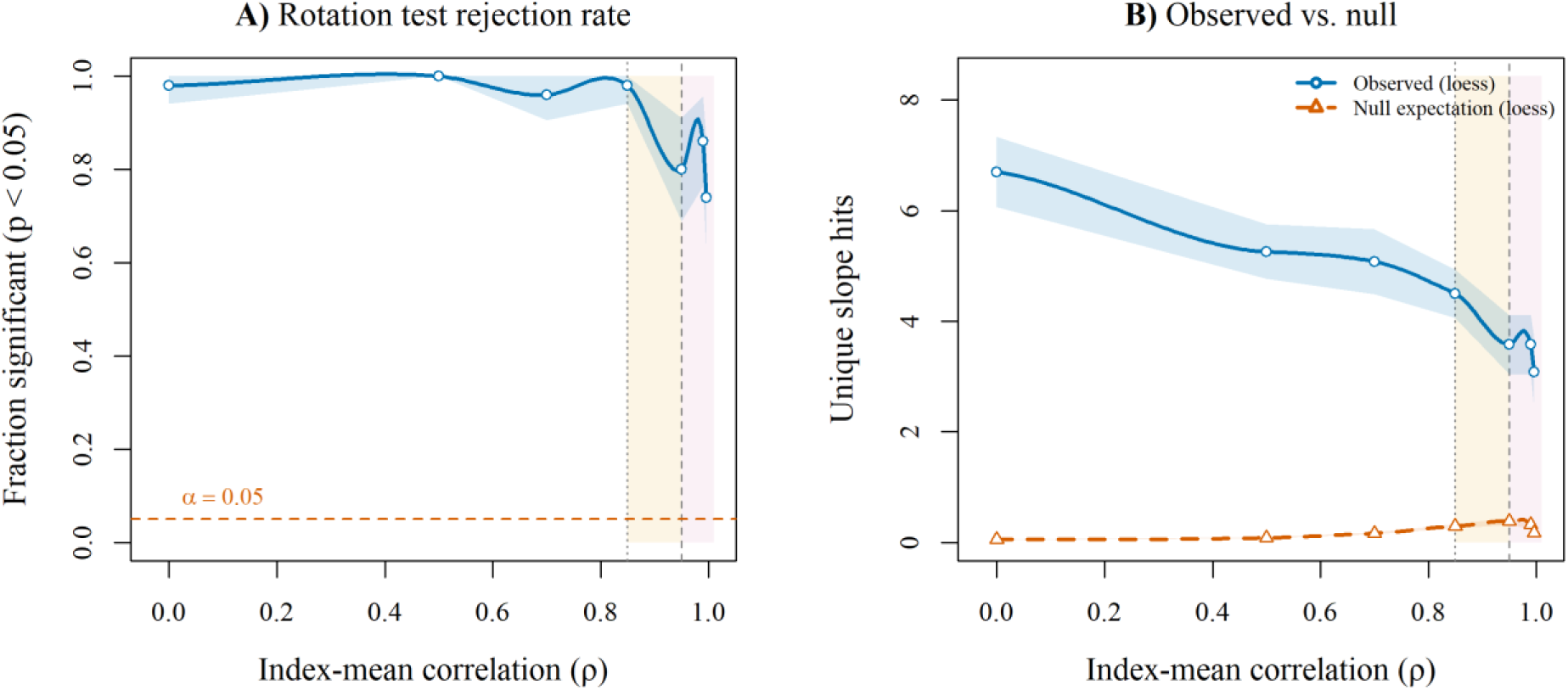
Rotation test discriminatory power (Experiment 12). (a) Rejection rate of the pure-DS null (*P* < 0.05), loess-smoothed with 95% CI bands. Orange dashed line: *α* = 0.05 nominal rate. (b) Observed unique slope QTL count (blue) versus null expectation (orange, dashed).

**Figure S6.**
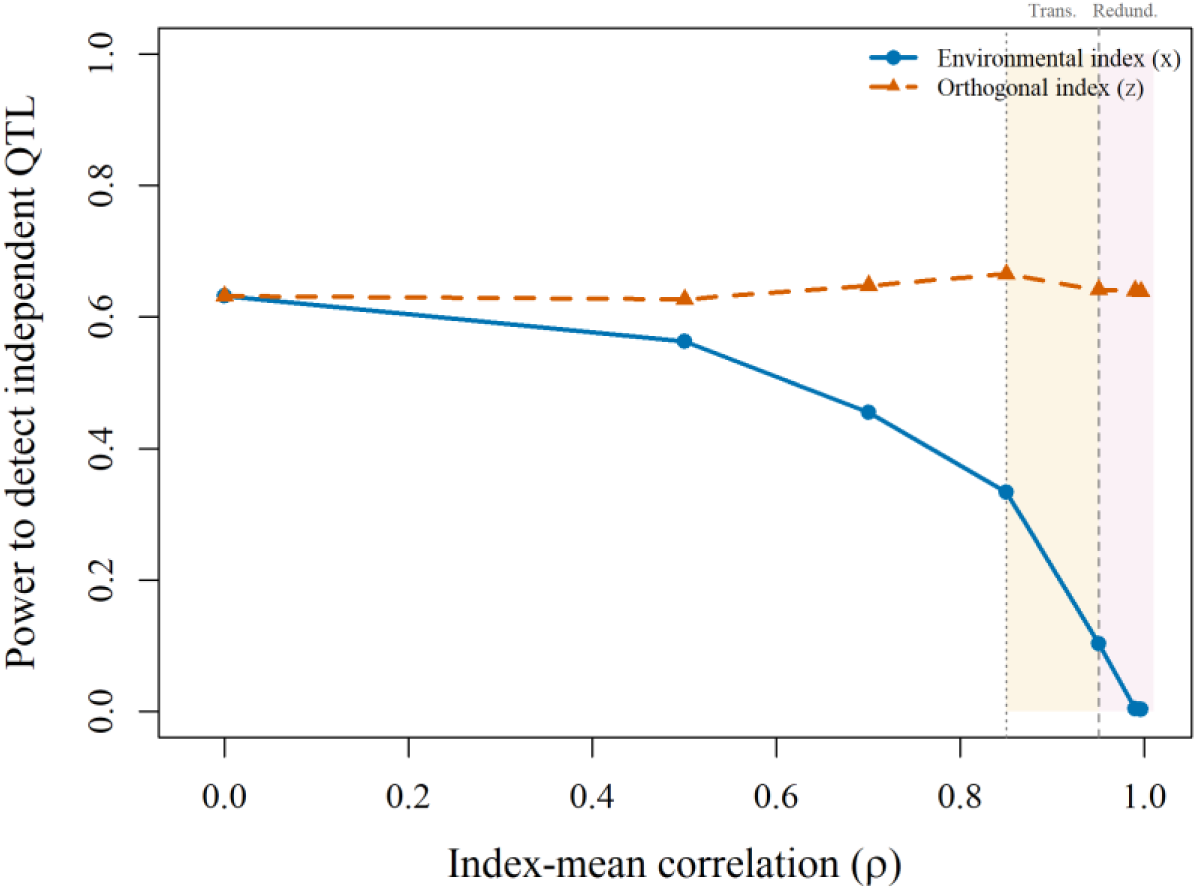
Multi-index versus single-index power (Experiment 13). Power to detect independent QTL for the primary index (blue, solid) and an orthogonal secondary index (orange, dashed) as a function of *ρ*. Shaded regime bands as in Figure 1.

**Figure S7.**
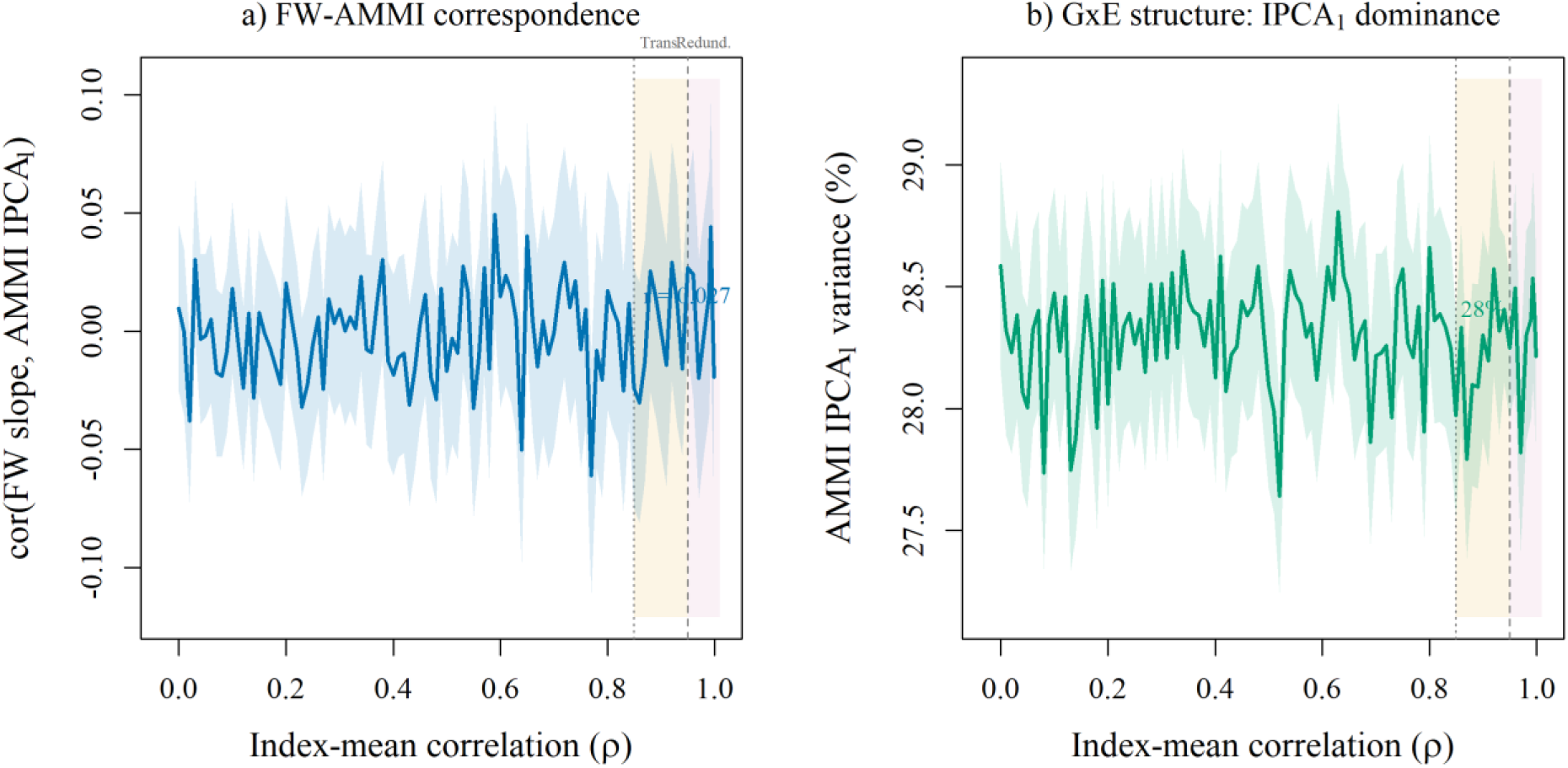
Finlay-Wilkinson and AMMI correspondence (Experiment 1). (a) Correlation between genotype-level FW slope and AMMI IPCA₁ score as a function of *ρ*, with 95% CI. (b) Percentage of G×E variance explained by IPCA₁.

**Figure S8.**
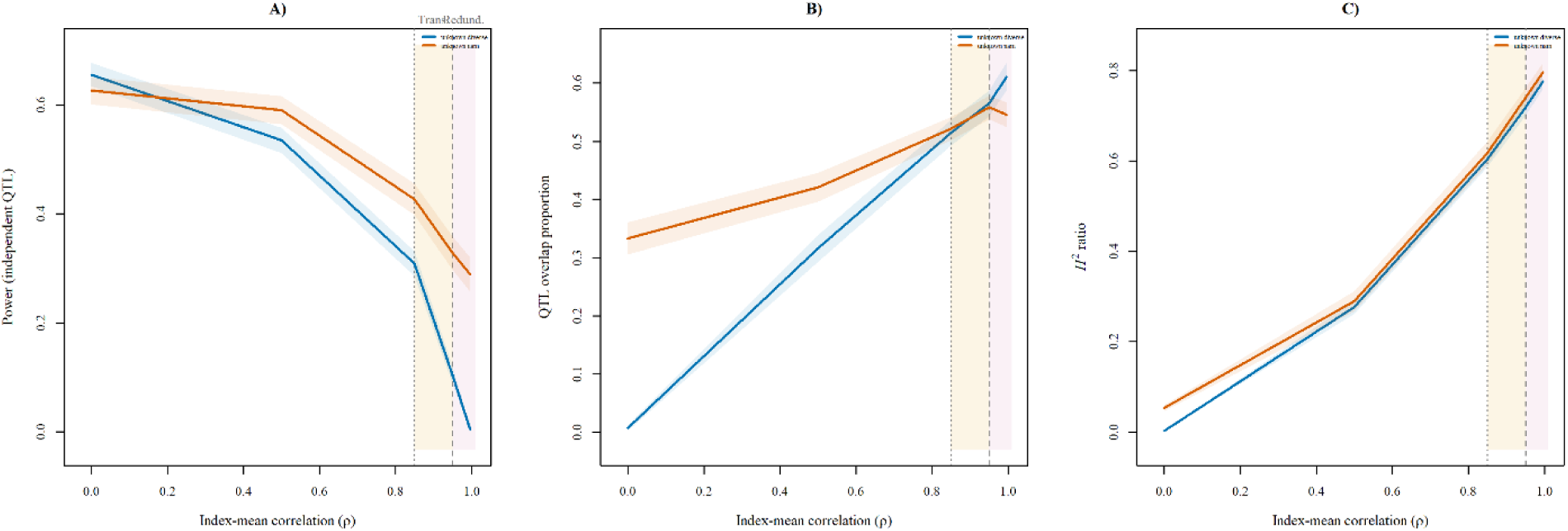
PCA-corrected LMM GWAS robustness (Experiment 18) for diversity panel (blue) and NAM (orange). (a) Power to detect independent QTL. (b) QTL overlap proportion. (c) Heritability ratio (H²slope / H²intercept).

**Figure S9.**
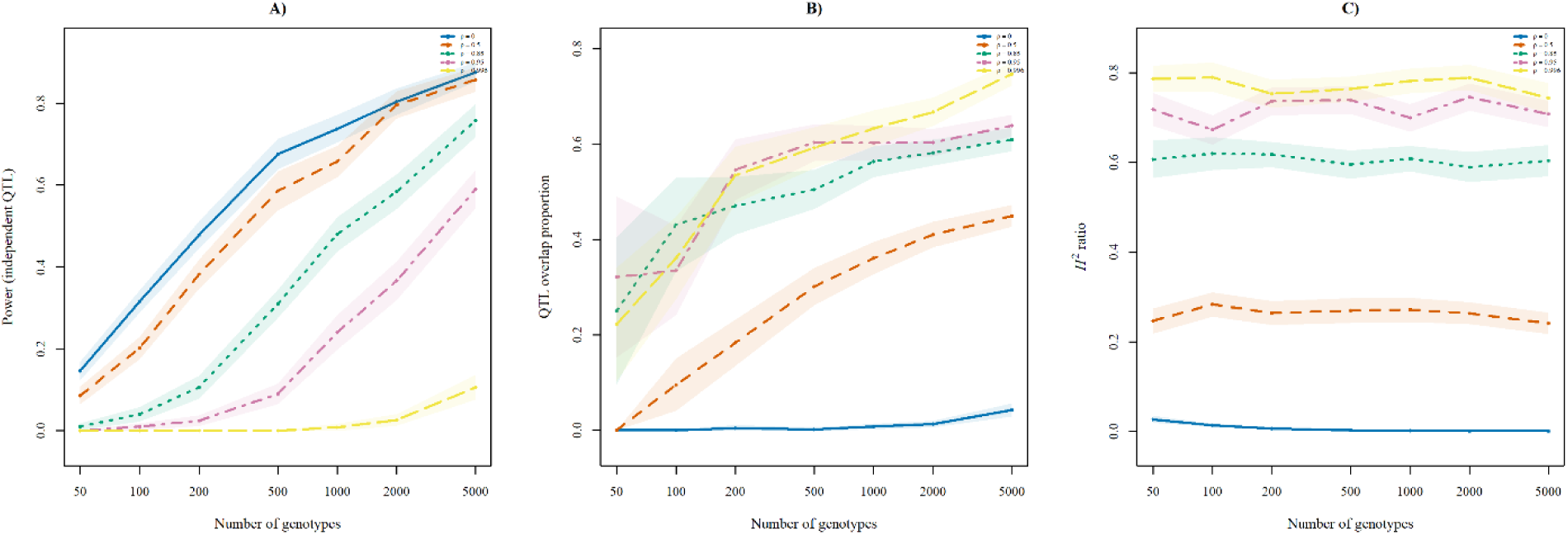
Sample size sensitivity (Experiment 9; *N* = 50–5,000) across five *ρ* levels. (a) Power to detect independent QTL. (b) QTL overlap proportion. (c) Heritability ratio.

**Figure S10.**
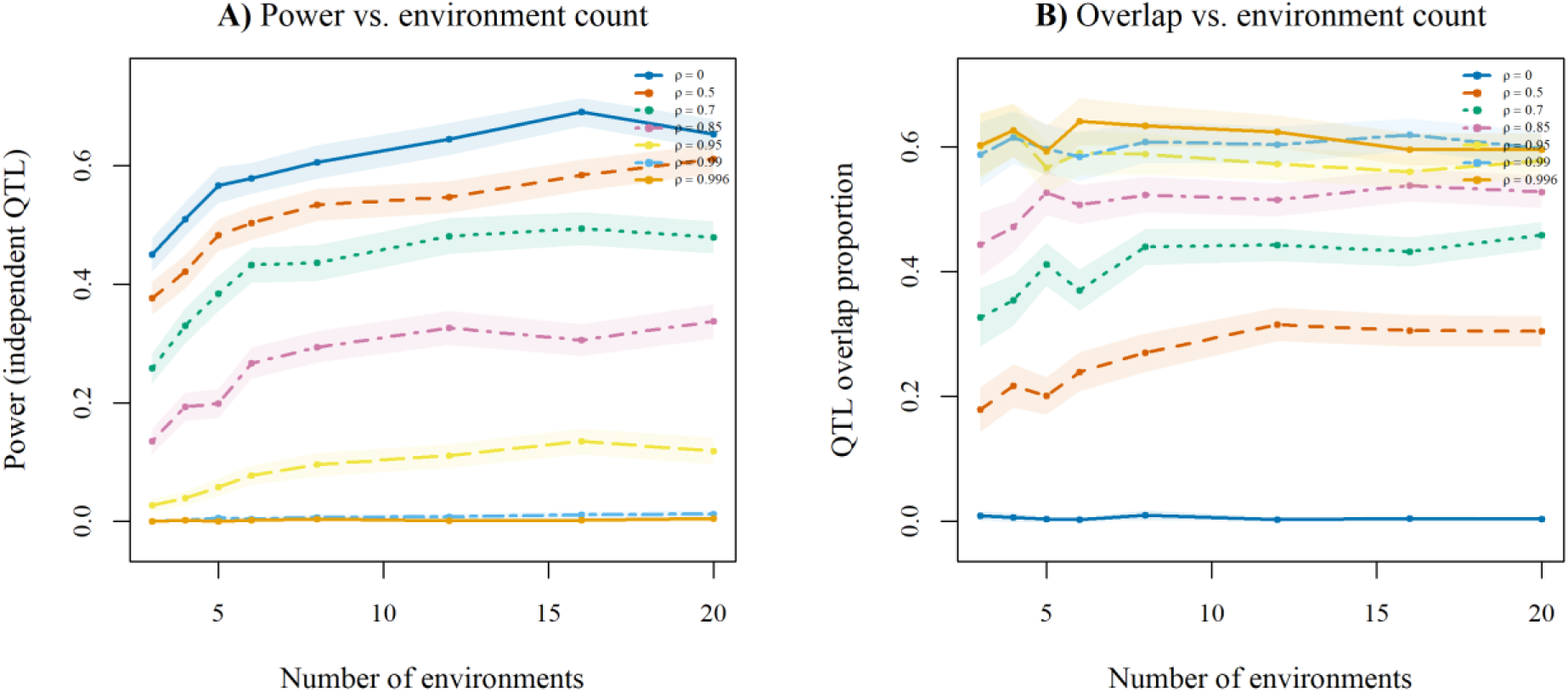
Environment count sensitivity (Experiment 6; 3–20 environments) across seven *ρ* levels. (a) Power to detect independent QTL. (b) QTL overlap proportion.

**Supplemental Table S1.** Complete cross-study meta-analysis dataset (47 entries, 27 publications, 11 species) with all extracted parameters. Available as cross_study.csv in the online supplement.

**Supplemental Table S2.** Comparison of simulation predictions to TC24 empirical values at matched *ρ*. Emp.cor = empirical −log₁₀(*P*) correlation. Sim.cor = bootstrap mean simulation prediction. Ratio = empirical/simulation. Reg. = Regime: C = Constrained (≥ 0.99), T = Transitional (0.85–0.99). N_1_ = intercept markers; N_2_ = slope markers. Ten of the 19 traits of Tibbs-Cortes et al. (2024) are shown, those for which ρ is reported or computable

| Trait | $\rho$ | N1 | N2 | Jaccard | Emp.cor | Sim.cor | Ratio | Reg. |
| --- | --- | --- | --- | --- | --- | --- | --- | --- |
| DTA | 0.999 | 2810 | 19828 | 0.004 | 0.11 | 0.19 | 0.58 | C |
| DTS | 0.998 | 3399 | 18994 | 0.066 | 0.10 | 0.19 | 0.53 | C |
| ASI | 0.996 | 757 | 2393 | 0.008 | 0.04 | 0.19 | 0.21 | C |
| EH | 0.993 | 1068 | 3299 | 0.013 | 0.09 | 0.18 | 0.50 | C |
| PH | 0.988 | 2095 | 4753 | 0.020 | 0.09 | 0.18 | 0.50 | T |
| LL | 0.960 | 481 | 139 | 0.000 | 0.02 | 0.17 | 0.12 | T |
| LW | 0.980 | 677 | 203 | 0.005 | 0.07 | 0.18 | 0.39 | T |
| TPBN | 0.975 | 206 | 169 | 0.013 | 0.21 | 0.18 | 1.17 | T |
| TKW | 0.999 | 97 | 1090 | 0.009 | 0.19 | 0.19 | 1.00 | C |
| T20KW | 0.998 | 200 | 1024 | 0.000 | 0.08 | 0.19 | 0.42 | C |

